# A dual-mechanism antibiotic targets Gram-negative bacteria and avoids drug resistance

**DOI:** 10.1101/2020.03.12.984229

**Authors:** James K. Martin, Maxwell Z. Wilson, Gabriel M. Moore, Joseph P. Sheehan, André Mateus, Sophia Hsin-Jung Li, Benjamin P. Bratton, Hahn Kim, Joshua D. Rabinowitz, Athanasios Typas, Mikhail M. Savitski, Zemer Gitai

**Author notes:** Please address correspondence to Zemer Gitai.

## Abstract

The rise of antibiotic resistance and declining discovery of new antibiotics have created a global health crisis. Of particular concern, no new antibiotic classes have been approved for treating Gram-negative pathogens in decades. Here, we characterize a compound, SCH-79797, that kills both Gram-negative and Gram-positive bacteria through a unique dual-targeting mechanism of action (MoA) with undetectably low resistance frequencies. In an animal host model, SCH-79797 reduces pathogenesis of *Acinetobacter baumannii*, a drug-resistant Gram-negative pathogen. To characterize the MoA of SCH-79797 we combined quantitative imaging, proteomic, genetic, metabolomic, and cell-based assays. This pipeline shows that SCH-79797 has two independent cellular targets, folate metabolism and bacterial membrane integrity, and outperforms combination treatments with other antifolates and membrane disruptors in killing MRSA persisters. Thus, SCH-79797 represents a promising lead antibiotic and suggests that combining multiple MoAs onto a single chemical scaffold may be an underappreciated approach to target challenging bacterial pathogens.

## INTRODUCTION

More than twenty unique classes of antibiotics were characterized in the 30 years following the discovery of penicillin in 1929 (Coates et al., 2011; Davies, 2006). However, a combination of scientific and economic factors have slowed the discovery and development of these life-saving molecules to the extent that only six new classes of antibiotics have been approved in the past 20 years, none of which are active against Gram-negative bacteria (Butler et al., 2016). This decline in the discovery of new antibiotic classes, coupled with the evolution of multi-drug resistant bacteria and horizontal transfer of resistance mechanisms, has created a public health crisis that is predicted to only escalate in the coming years (Culyba et al., 2015; Hofer, 2018; O’Neill, 2014).

Recent efforts have begun to reinvigorate antibiotics research, but most of this work has resulted in compounds that function via similar mechanisms to those of traditional antibiotics. For example, finafloxacin, an exciting new fluoroquinolone antibiotic that was recently approved to treat ear infections caused by *Pseudomonas aeruginosa*, is more effective than other fluoroquinolones because it maintains its potency in acidic environments (McKeage, 2015). However, finafloxacin is still susceptible to the same resistance mechanisms that affect other fluoroquinolones (Randall et al., 2016). The recent discovery of the natural product teixobactin suggests that it is possible to find compounds that selectively kill bacteria without being prone to resistance (Ling et al., 2015). However, teixobactin is only functional against Gram-positive bacteria and is a large molecule that is difficult to synthesize at commercial scales. Thus, there is still a strong need for characterizing new classes of antibiotics with distinct mechanisms of action (MoA).

An ideal antibiotic would be hard to develop resistance against, able to target Gram-negative bacteria, and easy to synthesize. It is important to note that while antibiotics that are not prone to resistance are attractive clinically, selecting for resistant mutants is the most common method for characterizing MoA, making the characterization of new antibiotic MoAs without resistance mutants a significant challenge. Phenotypic methods, such as macromolecular synthesis assays, have been previously used in such cases, as was done for teixobactin (Ling et al., 2015). However, these assays only allow the classification of molecules with previously-described MoAs (King and Wu, 2009). Thus, there is also a need for resistance-independent approaches for the *de novo* characterization of antibiotic MoA.

Here, we describe a compound, SCH-79797, that is bactericidal towards both Gram-negative and Gram-positive bacteria, including clinically important bacteria such *as Staphylococcus aureus* MRSA and *Acinetobacter baumannii*, with no signs of resistance. In an animal host model, SCH-79797 blocked infection by *A. baumannii* with low toxicity to the host at the dose required for effective antibiotic activity. To rapidly and efficiently classify the MoA of SCH-79797, we used a variant of a recently described quantitative imaging-based approach known as bacterial cytological profiling (BCP) (Nonejuie et al., 2013). This effort showed that SCH-79797 functions through a mechanism distinct from that of any known class of antibiotics. In the absence of being able to evolve resistance mutants, we used thermal proteome profiling (Savitski et al., 2014), CRISPRi genetic sensitivity (Peters et al., 2016), and metabolomic profiling (Kwon et al., 2008, 2010) to characterize the MoA of SCH-79797. Using this multi-dimensional, systems-level approach, we identified the candidate targets of SCH-79797 as dihydrofolate reductase and the bacterial membrane. Classical enzymology and membrane permeability and polarization assays confirmed the targets identified by our high-throughput approaches. Finally, we used a derivative of SCH-79797 to demonstrate that the two activities of this compound can be separated. Thus, our findings identify and characterize a promising new antibiotic and provide a potential roadmap for future antibiotic discovery efforts.

## RESULTS

### SCH-79797 is a broad-spectrum, bactericidal antibiotic

With the aim of finding antibiotics with novel mechanisms of action (MoA), we began with an unbiased, whole-cell screening approach. To include antibiotics that target both Gram-negative and Gram-positive bacteria, we screened for compounds that inhibited the growth of *E. coli lptD4213*, which has a compromised outer membrane that makes it partially-permeable to antibiotics that would otherwise have difficulty penetrating the Gram-negative lipopolysaccharide (Ruiz et al., 2006). We screened a drug library of ∼33,000 unique compounds and one of our most potent hits was SCH-79797, a compound that had been previously reported as a human PAR-1 antagonist (Ahn et al., 2000). This finding was surprising since there are no PAR-1 homologs in bacteria. A recent report suggested that SCH-79797 increases the ability of neutrophils to kill bacteria, perhaps by directly functioning as an antibiotic (Gupta et al., 2018). Given that studies focusing on characterizing its anticoagulant activities suggested that at least 5 mg/kg SCH-79797 can be safely tolerated in animals (Gobbetti et al., 2012; Strande et al., 2007) and its emergence as a potential antimicrobial with no known bacterial target (Gupta et al., 2018), we decided to further characterize SCH-79797 as a candidate antibiotic.

To assess the spectrum of bacterial species susceptible to SCH-79797, we measured the minimal inhibitory concentration (MIC) of SCH-79797 against several clinically-relevant pathogens. In this study, we define MIC as the concentration of drug that results in no visible bacterial growth after 14h of growth at 37°C. We found that SCH-79797 significantly hindered the growth of multiple Gram-negative and Gram-positive pathogens such as *Neisseria gonorrhoeae*, two clinical isolates of *Acinetobacter baumannii*, and methicillin-resistant *Staphylococcus aureus* (MRSA) (Figure 1A). Using the *E. coli lptD4213* strain from our original screen, we found that SCH-79797 exhibits potent and rapid bactericidal activity against *E. coli lptD4213* (Figure 1B and C). SCH-79797 also exhibited similar bactericidal activity against a clinical isolate of *S. aureus* MRSA (USA300) (Tenover and Goering, 2009) suggesting that its bactericidal activity is not species-specific (Figure S1).

**Figure 1.**
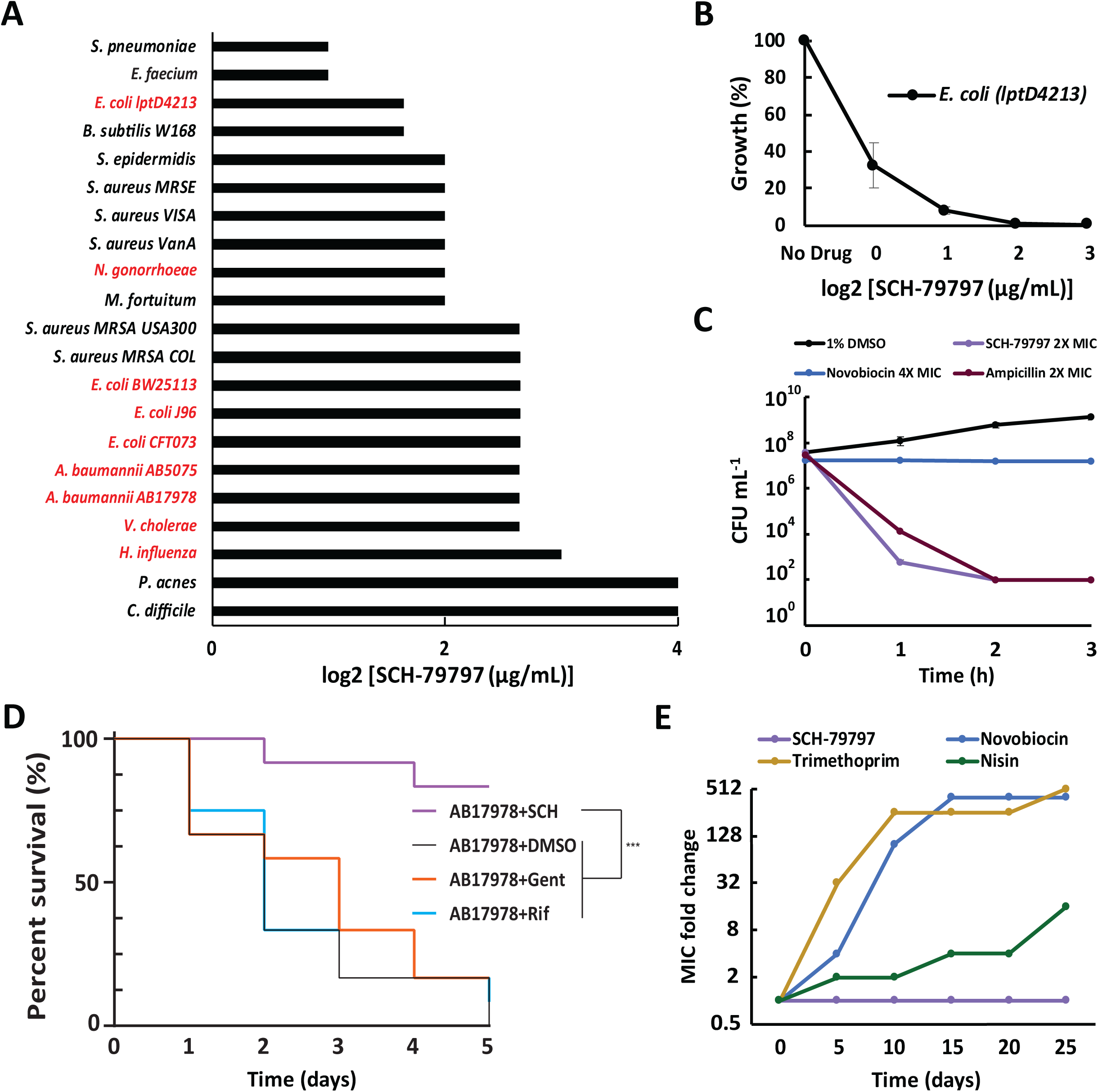
SCH-79797 is a broad-spectrum bactericidal antibiotic that is effective in an animal model and has a low frequency of resistance. A. The MIC of SCH-79797 against Gram-negative (red) and Gram-positive (black) bacteria. MIC here and in subsequent figures is defined as the concentration of drug that resulted in no visible bacterial growth. Bacteria were grown for 14h at 37°C and growth media for each bacterium is specified in table S1. B. The relative growth of *E. coli lptD4213* after treatment with SCH-79797. Bacterial growth was measured for 14h and the final optical density at 600nm (OD600) was plotted against drug concentration. Each data point represents 2 independent replicates. Mean ± s.d. are shown. C. Colony forming units (CFU ml^−1^) after 3-hour treatment of *E. coli lptD4213* with 1% DMSO, 2X MIC SCH-79797, 2X MIC ampicillin, and 4X MIC novobiocin. Data points at 1 × 10^2^ CFU ml^−1^ are below the level of detection. Each data point represents 3 independent samples and 3 technical replicates. Mean ± s.d. are shown. D. The percent survival of *A. baumannii* infected *G. mellonella* larvae after treatment with 2µl/larva of 100% DMSO, 67µg/larva SCH-79797, 6µg/larva gentamicin, and 67µg/larva rifampicin. Data represents a typical cohort (n = 12) from a biological triplicate. Mantel-cox statistics for the cohort were calculated with PRISM, and the pooled results are presented in the supplemental material (Figure S2C). E. Fold increase in resistance of *S. aureus* MRSA USA300 to SCH-79797, novobiocin, trimethoprim, and nisin after 25 days of serial passaging in 0.5X MIC of each drug and plotted on a log2 scale. Resistance was confirmed by remeasuring MIC’s from aliquots of each passage that were collected and stored at −80°C. Data represents one biological replicate and the data for the second replicate is shown in Figure S3A.

### SCH-79797 is effective in vivo and has a low frequency of resistance

Given SCH-79797’s promising ability to kill bacteria, we sought to determine if it can function as an effective antibiotic *in vivo*. To test its antibiotic activity in the context of an animal host infection, we focused on *A. baumannii* as it has emerged as an important Gram-negative pathogen that is targeted by relatively few available antibiotics, and has a well-established host animal model in the wax worm, *Galleria mellonella* (Gebhardt et al., 2015; Peleg et al., 2009). We first established that injecting *G. mellonella* with SCH-79797 at concentrations four times higher than the MIC of SCH-79797 towards *A. baumannii* did not result in higher host toxicity than the established antibiotics, gentamicin and rifampicin (Fig. S2A-B). We next tested the ability of SCH-79797 to treat infection of *G. mellonella* with a lethal dose of *A. baumannii* AB17978. Treatment with SCH-79797 significantly prolonged the survival of *A. baumannii*-infected *G. mellonella* (*P* < 0.001) (Figure 1D and S1C-D). The survival rate of *G. mellonella* treated with SCH-79797 exceeded that of gentamicin and rifampicin (Fig. 1D and S2C-D), which are standard antibiotics used to treat *A. baumannii* infections (Karlowsky et al., 2003; Viehman et al., 2014).

To further characterize the promise of SCH-79797 as an antibiotic, we attempted to determine the frequency with which bacteria develop resistance towards SCH-79797. Because spontaneous suppressors can restore *E. coli lptD4213*’s membrane barrier functionality, we focused our resistance studies on *S. aureus* MRSA (USA300) (Tenover and Goering, 2009). We were unable to isolate stable SCH-79797-resistant mutants upon plating ∼10^8^ CFU of MRSA USA300 onto agar containing 4X MIC of SCH-79797. We were also unable to isolate SCH-79797-resistant mutants upon plating ∼10^8^ CFU of *B. subtilis*, suggesting that the difficulty in developing resistant mutants is not species-specific. To address resistance rates more quantitatively, we serial-passaged 2 biologically independent cultures of *S. aureus* MRSA USA300 in 0.5X MIC of SCH-79797, as well as three control antibiotics: novobiocin, trimethoprim, and nisin. Over the course of 25 days, we successfully isolated mutants resistant to all the control antibiotics while no SCH-79797-resistant mutants emerged (Figure 1E, S3A). For novobiocin, trimethoprim, and nisin, resistance gradually increased throughout the experiment, but the resistance level remained constant for SCH-79797 (Figure 1E, S3A), indicating that these bacteria did not even acquire partial resistance to SCH-79797. To extend these findings to a Gram-negative species, we repeated our serial passaging study with 2 biologically independent cultures of *A. baumannii* AB17978 (Figure S3B). This study also found that *A. baumannii* resistance stayed constant for SCH-79797 but increased for all other antibiotics, supporting the conclusion that the lack of resistance to SCH-79797 is not species-specific.

### A variant of Bacterial Cytological Profiling suggests that SCH-79797 has a unique MoA

The inability to isolate SCH-79797-resistant mutants makes SCH-79797 an appealing candidate antibiotic but poses a challenge for determining its MoA. As a result, we used a quantitative imaging-based approach to determine if the MoA of SCH-79797 is similar to that of any previously-characterized antibiotics. Specifically, we modified a single-cell, high-content imaging methodology, known as Bacterial Cytological Profiling (BCP) (Nonejuie et al., 2013). The logic of BCP is that antibiotics with similar MoA result in similar death phenotypes such that by quantifying how bacteria appear upon death, we can gain insight into the cause of death (much like a bacterial autopsy). Here, we applied our BCP analysis to a training set of 37 distinct antibiotics with known MoA as well as to SCH-79797. For each compound, we treated *E. coli lptD4213* with 5X MIC of an antibiotic for 2 hours, stained with three dyes that report on nucleoid morphology (DAPI), membrane morphology (FM4-64), and membrane integrity (SYTOX Green), and imaged the cells at high resolution (Figure 2A). For each condition we imaged ∼100 cells and quantified 14 parameters reflecting various morphological and fluorescence features (Supplementary Table 2).

**Figure 2.**
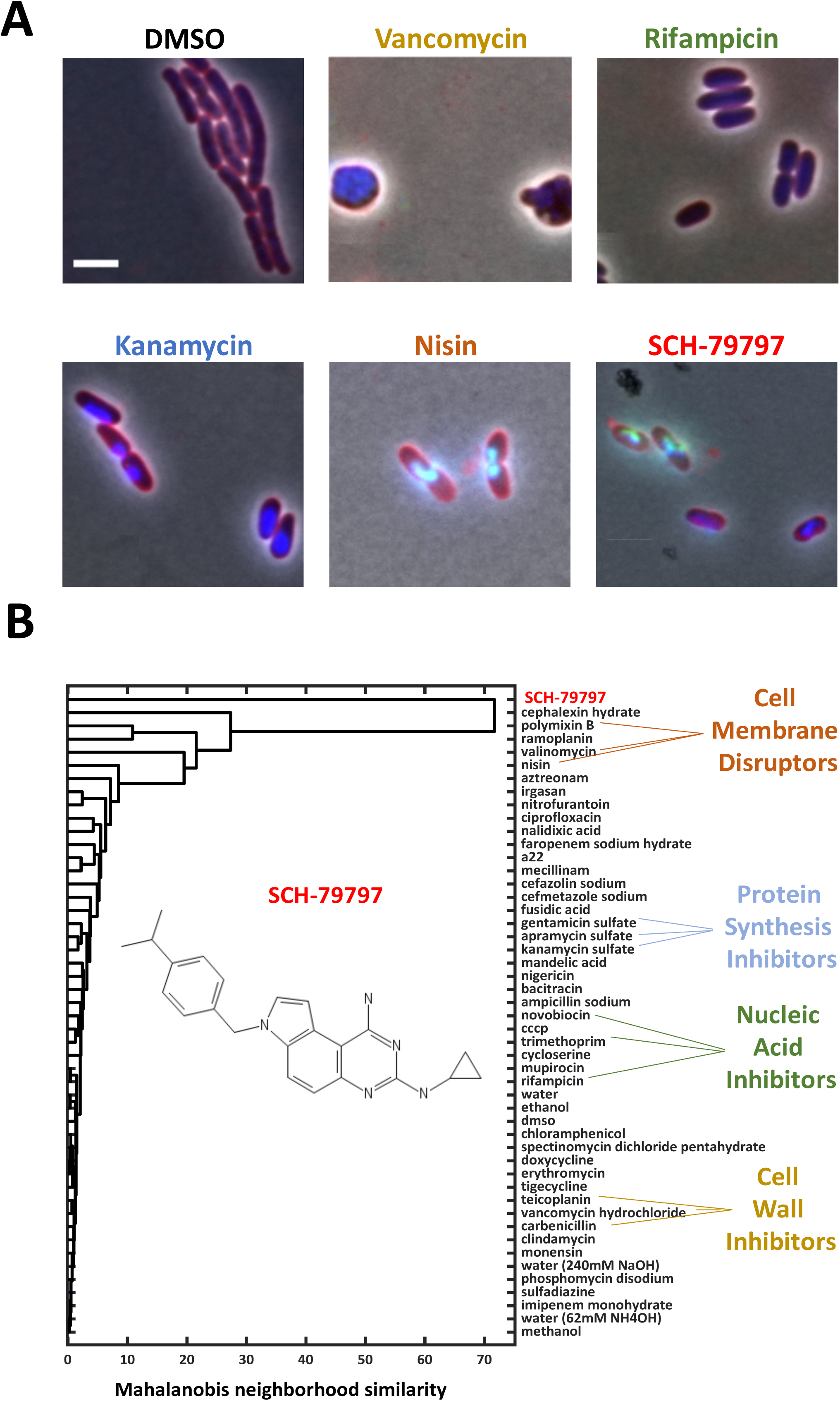
Bacterial Cytological Profiling indicates that SCH-79797 functions by a mechanism distinct from known classes of antibiotics. A. Fluorescent images of *E. coli lptD4213* cells treated with antibiotics representative of 5 different antibiotic classes. Cells were treated for 2h with 5X MIC of each drug. Merged image channels are phase (grey), FM4-64 (red), Dapi (blue), and SYTOX (green). All images are at the same magnification and the scale bar is 1μm. B. Comparison of cytological profiles of known antibiotics with the cytological profile of SCH-79797. Single-linkage clustered vectors of Mahalanobis distances from each antibiotic treatment group were compared to that of all other antibiotic treatment groups. Linkage is included in the dendrogram.

Since we had gold standards of the BCP results of SCH-79797 and of several antibiotics representing different classes and sub-groups within classes, we applied a machine learning approach to classify the BCP data. For each treatment group, we populated a ‘neighborhood representation’ vector with the one-way Mahalanobis distance as measured from the single-cell feature mean in question, to the distribution measurement of all other treatment groups. This distance is normalized by the covariance matrix of the antibiotic treatment group so that dimensions with large amounts of variance are deemed closer, while distances of dimensions with less variance are considered farther away. We then used single-linkage clustering to cluster treatment groups by their neighborhood representation vectors, such that samples whose neighborhoods were similar would be clustered together. This analysis indicated that SCH-79797 resulted in a phenotypic death-state that was different from the other antibiotics tested (Figure 2B), suggesting that SCH-79797 possesses a MoA distinct from that of any of the antibiotics in our training set.

### Thermal profiling and CRISPRi genetics demonstrate that SCH-79797 targets dihydrofolate reductase (DHFR)

In the absence of resistant mutants or similarity to antibiotics with known MoA by BCP, we turned to a high-throughput proteomics-based approach for *de novo* identification of candidate SCH-79797 targets. Specifically, we used thermal proteome profiling, an assay that uses mass spectrometry to compare the thermal stability of the entire proteome with and without drug treatment (schematized in Figure 3A) (Mateus et al., 2018; Savitski et al., 2014). Briefly, intact cells or cell lysate samples treated with a range of compound concentrations are heated to a series of increasing temperatures and the soluble proteins at each temperature are collected (Becher et al., 2016). Proteins that bind to the drug are thermally stabilized, which leads to a shift in the temperature at which those proteins precipitate (Figure 3A). Using *E. coli lptD4213*, we treated intact cells and cell lysates with SCH-79797 and found that it significantly shifted the thermal stability of dihydrofolate reductase (DHFR) (Figure 3B and S4A). The fact that the same result was observed with both intact cells and cell lysates (Figure 3B and S4A) suggests that SCH-79797 enters *E. coli* cells and directly binds to *E. coli* DHFR. As a positive control, we used a well-characterized antibiotic that targets DHFR, trimethoprim, and found that it also thermally stabilizes its known target, DHFR (Figure 3C and S4B).

**Figure 3.**
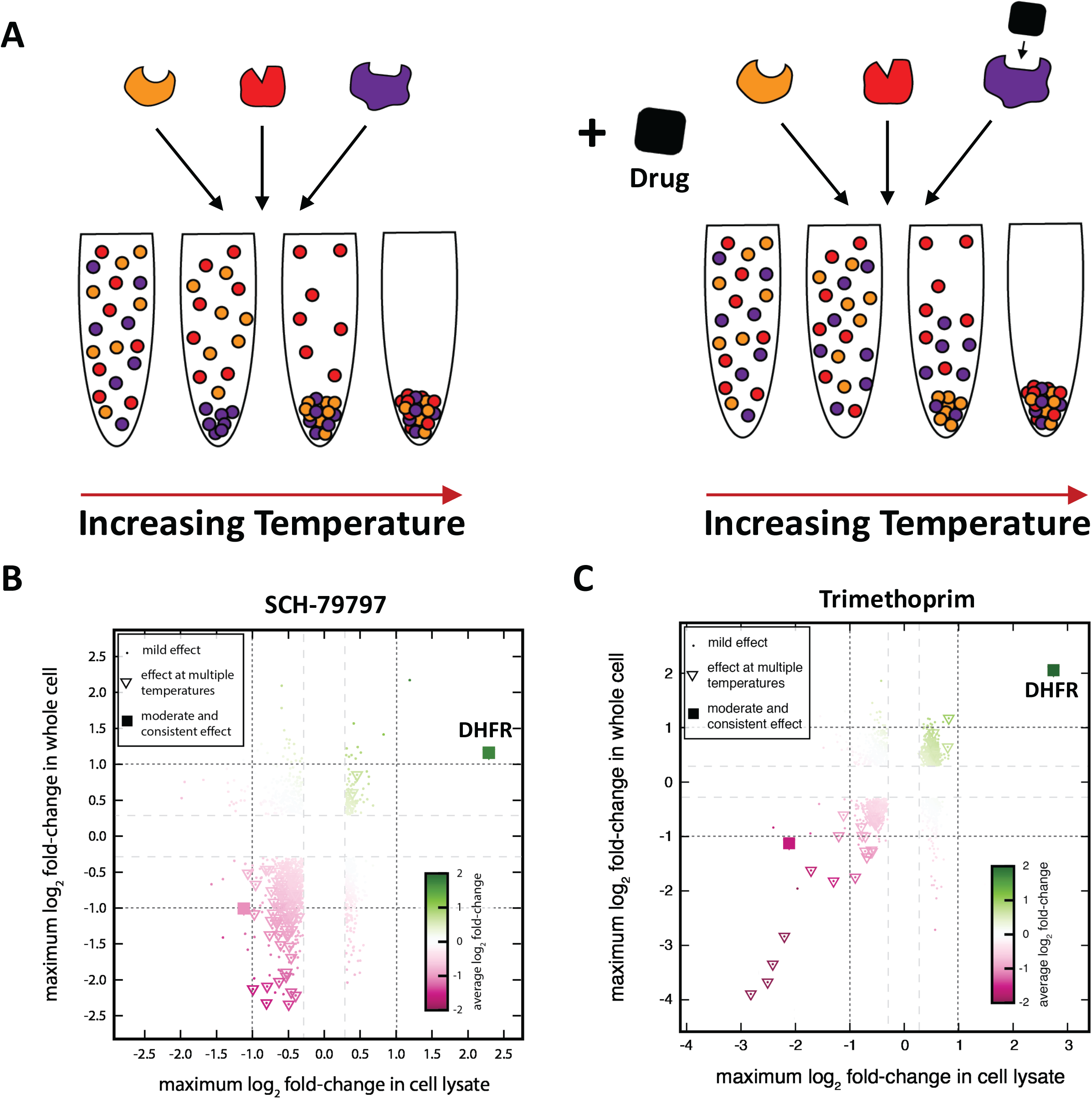
Thermal proteome profiling suggests that SCH-79797 binds DHFR. A. Schematic of the thermal shift assay that compares the thermal stability of the entire proteome with and without drug treatment. Protein samples are aliquoted, and each aliquot is heated to an increasing temperature. The relative fraction of soluble and insoluble proteins is then determined for each aliquot by ultracentrifugation and mass spectrometry. B-C. The relative thermal stability of the soluble *E. coli lptD4213* proteome after treatment of whole cell and cell lysate samples with SCH-79797 and trimethoprim. Changes in thermal stability were determined by measuring changes in the abundance of soluble protein across 10 different temperatures ranging from 42-72°C and 4 drug concentrations and a vehicle control. For each point, the color indicates the maximal effect size across all temperatures and the largest change in abundance across all concentrations. Squares represent the proteins with a change in abundance of at least 25% at three or more temperatures. To be considered consistent, the change in abundance of a protein had to show the same sign at least 90% of the time and have an effect size of at least 2-fold in either whole cells or cell lysates. Triangles represent a milder effect where at least one temperature had a change in abundance of at least 25% in both whole cell and cell lysate treatments.

To test both the physiological significance and species-specificity of the suggestion that SCH-79797 binds to DHFR, we took advantage of a collection of *B. subtilis* essential gene CRISPR-interference (CRISPRi) knockdown mutants (Peters et al., 2016). In each of these mutants, an essential gene is targeted by CRISPRi to reduce its expression ∼3-fold. A strain with reduced levels of the SCH-79797 target should be sensitized to sub-lethal doses of SCH-79797. Given the thermal profiling result, we focused on mutants in the folate biosynthesis pathway (Figure 4A). As a negative control, we confirmed that CRISPRi knockdowns of genes unrelated to folate metabolism are not sensitized to SCH-79797 (Figure S5). As a positive control for our assay, we used trimethoprim. We confirmed that dihydrofolate reductase (*dfrA*) and dihydrofolate synthase (*folC*) (an enzyme that acts upstream of DfrA) knockdowns are hypersensitive to trimethoprim, while enzymes that function downstream of DfrA, *folD* and *glyA*, are not (Figure 4B). SCH-79797 exhibited the same genetic sensitivity pattern as trimethoprim in that both *dfrA* and *folC*, but not *folD* and *glyA* knockdowns, were sensitized to SCH-79797 (Figure 4B).

**Figure 4.**
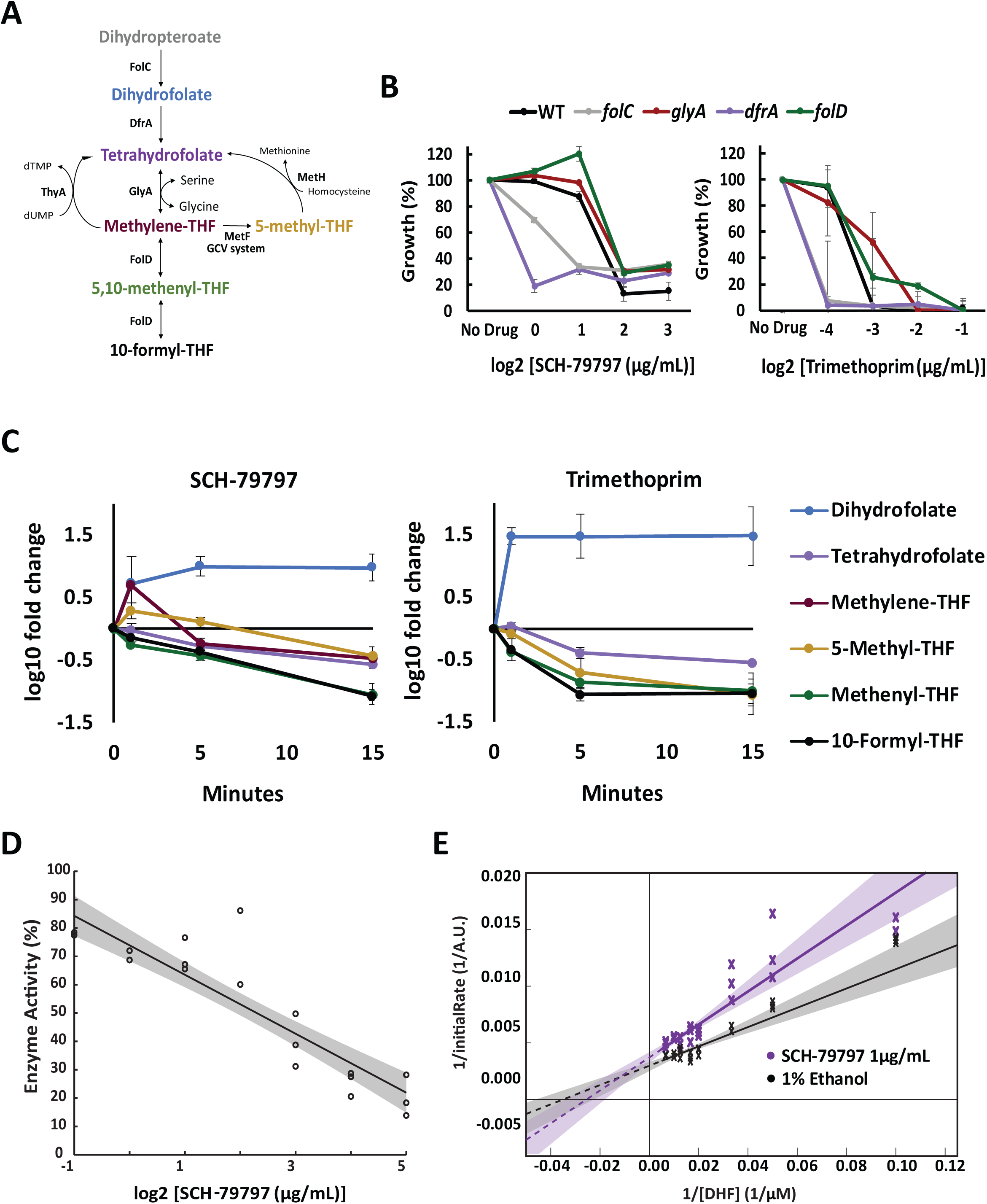
SCH-79797 targets folate metabolism by competitively inhibiting DHFR. A. A partial representation of the folate synthesis pathway. B. The growth of CRISPRi *B. subtilis* knockdown mutants involved in folate synthesis relative to a DMSO-treated control after SCH-79797 and trimethoprim treatment. Bacterial growth was measured for 14h and the final OD600 of each condition was plotted against drug concentration. Each data point represents 2 independent replicates. Mean ± s.d. are shown. C. Metabolomic analysis of *E. coli* NCM3722 cells treated with 0.5X MIC SCH-79797 or trimethoprim. Samples were taken 0, 5, 10, and 15 min. after drug treatment. Folate metabolite abundance at each time point was quantified relative to the DMSO-treated control samples at the initial timepoint. 3 independent replicates of this experiment were performed. Each data point represents 3 independent replicates. Mean ± s.d. are shown. D. The enzymatic activity of DHFR upon SCH-79797 treatment, relative to a vehicle-treated (ethanol) control. A linear-fit was applied to the resulting plot to determine the IC50 and 3 independent replicates are shown. The shaded region represents the 90% confidence interval of the fit. E. A Lineweaver-Burk plot of the enzymatic activity of DHFR after treatment with 1μg/mL SCH-79797 and a 1% ethanol control. Fits to the Michaelis-Menten equation are shown with shaded regions corresponding to 90% confidence intervals.

### SCH-79797 inhibits DHFR activity in cells and in vitro

To determine how SCH-79797 affects folate metabolism in living cells, we used mass spectrometry to measure the relative abundance of folate metabolite pools in *E. coli* NCM3722 treated with SCH-79797. *E. coli* NCM3722 was used because these bacteria lack mutations that disrupt primary metabolism in other lab strains of *E. coli* (Soupene et al., 2003). *E. coli* NCM3722 cells were grown in Gutnick Minimal Media and treated with 1X MIC SCH-79797 for 15 minutes (Kwon et al., 2008, 2010). In response to SCH-79797 treatment, the levels of the DHFR substrate, 7,8-dihydrofolate (DHF), rose approximately 10-fold compared to untreated cells, while the levels of folate metabolites downstream of DHF dropped significantly (Figure 4C). This metabolic response is characteristic of DHFR inhibition as we observed a similar pattern upon treatment with trimethoprim (Figure 4C).

To determine whether SCH-79797 inhibits DHFR *in vitro*, we obtained purified *E. coli* DHFR protein and measured its enzymatic activity in the presence of SCH-79797. We found that SCH-79797 has an IC50 of 2.5 ± 0.6 µg/mL against DHFR (Figure 4D). We also measured the initial velocity of DHFR activity at various DHF substrate concentrations in the presence or absence of SCH-79797 to establish if SCH-79797 acts competitively or non-competitively. Fitting our data to the Michaelis-Menten equation demonstrated that 1 µg/mL SCH-79797 increases the K_m_ from 29 ± 9 uM to 39 ± 13 uM and decreases the V_max_ of DHFR from 3300 ± 300 A.U. to 2700 ± 300 A.U.. These results indicate that SCH-79797 functions at least partially as a competitive inhibitor of DHFR’s activity on its DHF substrate (Figure 4E).

### SCH-79797 also disrupts bacterial membrane potential and permeability

The similarities between SCH-79797 and trimethoprim with respect to DHFR inhibition helped confirm DHFR as a target of SCH-79797 but were also surprising because these two compounds did not generate similar profiles in our BCP analysis (Figure 2B). One potential explanation is that SCH-79797 has additional targets that are not shared with trimethoprim. If this was the case, we would expect that cells resistant to trimethoprim would still be susceptible to SCH-79797. Previous studies demonstrated that resistance to trimethoprim can be achieved by deleting *thyA* and supplementing the media with thymine (Amyes and Smith, 1975). We confirmed that deleting *thyA* from *E. coli lptD4213* in the presence of excess thymine led to trimethoprim resistance (Figure 5A). However, these cells showed no change in their sensitivity to SCH-79797 (Figure 5A), suggesting that SCH-79797 is likely to have a second, folate-independent MoA.

**Figure 5.**
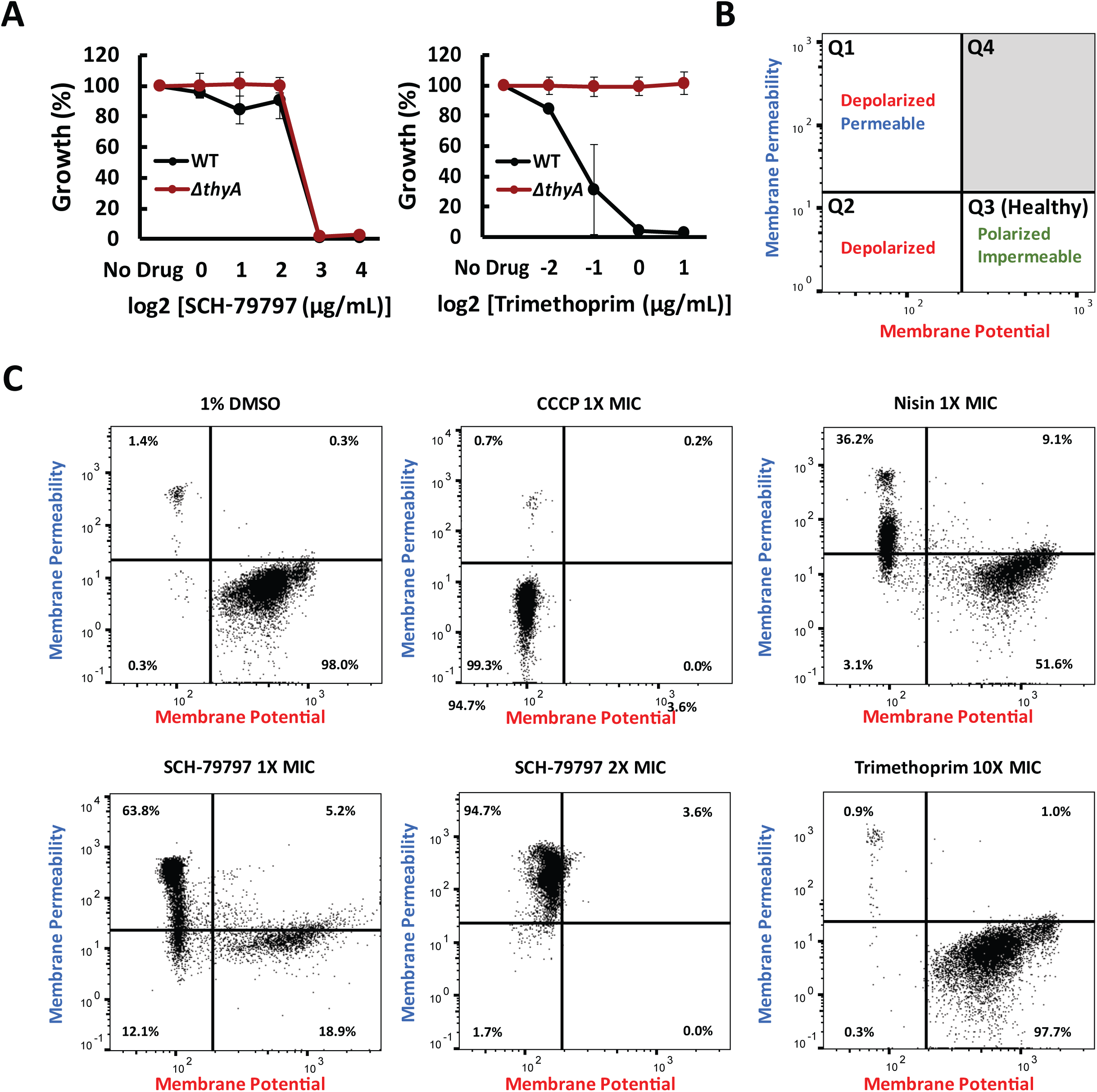
SCH-79797 is distinct from other DHFR inhibitors and disrupts membrane integrity. A. The growth of WT and *ΔthyA E. coli lptD4213* relative to a DMSO-treated control after SCH-79797 and trimethoprim treatment. Bacterial growth was measured for 14h and the final OD600 of each condition was plotted against drug concentration. Each data point represents 2 independent replicates. Mean ± s.d. are shown. B. Schematic of flow cytometry data showing the expected results for each class of polarized, depolarized, permeable and impermeable bacteria. C. Flow cytometry analysis of the membrane potential and permeability of *E. coli lptD4213* cells after 15 min. incubation with 1% DMSO, 1X MIC CCCP, 1X MIC nisin, 1X and 2X MIC SCH-79797, and 10X MIC trimethoprim. CCCP and nisin treatment served as depolarizing and permeabilizing controls respectively. These controls were used to define the quadrants outlined in (B).

To obtain clues about the potential additional MoA of SCH-79797, we revisited our fluorescent BCP images of *E. coli lptD4213* cells treated with SCH-79797. We observed SYTOX Green staining in some of the bacteria (Figure 2A), suggesting that SCH-79797 compromises the integrity of the bacterial membrane. To directly quantify the effect of SCH-79797 on bacterial membrane integrity, we used flow cytometry to measure the membrane potential and permeability of *E. coli lptD4213* in the presence of the fluorescent dyes, DIOC_2_(3) and TO-PRO-3. DIOC_2_(3) is a cationic dye that accumulates in the cytoplasm of cells with an active membrane potential and shifts its fluorescence from red to green in these cells, providing a measure of membrane potential (Figure 5B). TO-PRO-3 is a nucleic acid stain that only accumulates in cells with compromised membranes, providing an independent measure of membrane permeability (Figure 5B). As positive controls, we showed that CCCP, a membrane-decoupler that affects membrane potential but not permeability, and nisin, a pore-forming antibacterial peptide that disrupts both membrane potential and permeability, caused the expected shifts in both DIOC_2_(3) and TO-PRO-3 staining (Figure 5C). As negative controls, we confirmed that antibiotics that do not target the membrane, including ampicillin, rifampicin, and novobiocin, do not shift DIOC_2_(3) or TO-PRO-3 staining (Figure S6). After 15 minutes of treatment with SCH-79797 at MIC levels, DIOC_2_(3) and TO-PRO-3 staining revealed significant defects in both membrane polarization and permeability (Figure 5C). These effects on the membrane are not secondary consequences of DHFR inhibition, as trimethoprim-treated *E. coli* showed no significant changes in DIOC_2_(3) and TO-PRO-3 staining (Figure 5C). The membrane-targeting effect of SCH-79797 is also not species-specific, as similar results were seen with SCH-79797-treated *B. subtilis* W168 (Figure S7). These findings indicate that independent of its ability to inhibit DHFR activity, SCH-79797 disrupts both membrane potential and permeability.

### SCH-79797 treatment can kill bacteria in contexts where combination therapy fails

Having established that SCH-79797 disrupts both folate metabolism and membrane integrity, we sought to determine if these two targets can together explain how SCH-79797 kills bacteria. To address this question, we used BCP analysis to compare the cell morphology of bacteria treated with SCH-79797, to that of bacteria treated with trimethoprim and nisin, two different antibiotics that target DHFR and membrane integrity, respectively (Nonejuie et al., 2013; Wilson et al., 2016). Qualitative inspection suggested that when stained with DAPI, FM4-64, and SYTOX Green, SCH-79797 treated *E. coli* appeared similar to *E. coli lptD4213* cells treated with both trimethoprim and nisin (Figure 6A). Quantification of the images confirmed that SCH-79797 closely clusters with the co-treatment of trimethoprim and nisin (Figure 6A). The fact that SCH-79797 clusters more closely to the co-treatment than to the individual treatments with trimethoprim or nisin reinforces the conclusion that SCH-79797 kills bacteria by targeting both DHFR and the membrane. There are no other antibiotics that have been shown to target both folate metabolism and membrane integrity, indicating that SCH-79797 represents an antibiotic with a unique MoA. This result also explains why SCH-79797 failed to cluster with any of the known antibiotics in our BCP analysis (Figure 2B).

**Figure 6.**
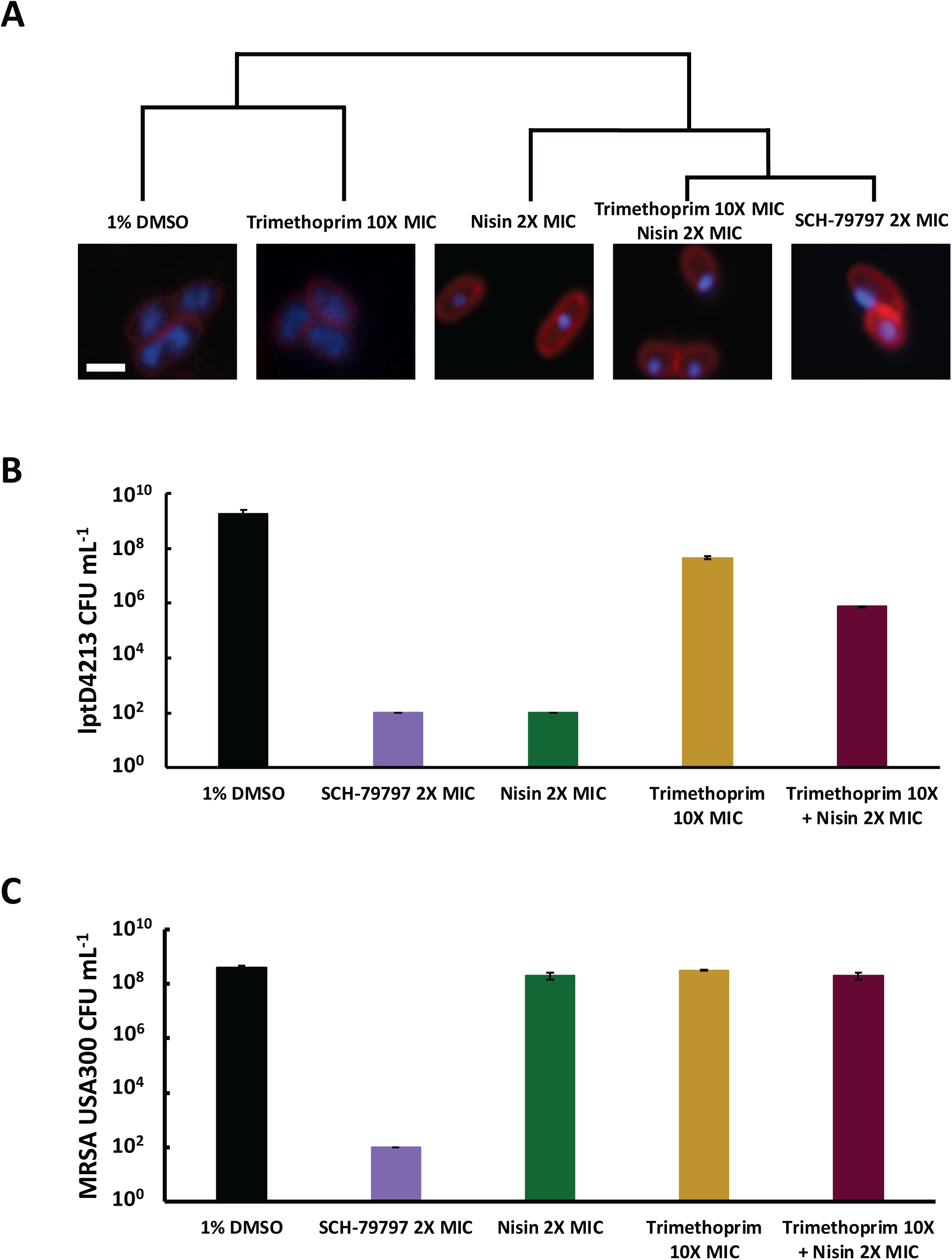
SCH-79797 mimics co-treatment with folate metabolism and membrane integrity disruptors but can be more effective than their combination. A. BCP analysis of *E. coli lptD4213* cells after 30 min. of treatment with 1% DMSO, 1X MIC SCH-79797, 10X MIC trimethoprim, 2X MIC nisin, and the combination of 10X MIC trimethoprim and 2X MIC nisin. Cytological profiles were clustered by the first three principal components that account for at least 90% of the variance between samples. Cells were stained with DAPI, FM4-64, and SYTOX Green. Shown here are the merged images of Dapi (blue) and FM4-64 (red) and the scale bar is 1μm. B-C. The viability of (B) *E. coli lptD4213* and (C) *S. aureus* MRSA USA300 persister cells measured in CFU mL^−1^ after 2 hours of treatment with 1% DMSO, 1X MIC SCH-79797, 10X MIC trimethoprim, 2X MIC nisin, and the combination of 10X MIC trimethoprim and 2X MIC nisin. Each bar represents 3 independent samples and 3 technical replicates. Mean ± s.d. are shown.

Our findings that SCH-79797 has the same MoA as combined trimethoprim and nisin treatment raised the question of whether there is a benefit to combining two targeting mechanisms onto a single molecule. Combination antibiotic therapy has been suggested as a potential means of circumventing the rise of antibiotic resistance (Tamma et al., 2012; Tyers and Wright, 2019) but it has remained unclear whether it is better to combine multiple activities on the same molecule. To probe this issue, we measured the synergy of trimethoprim and nisin against *E. coli lptD4213* and MRSA USA300 persister cells and compared their combined effectiveness to that of SCH-79797. Interestingly, when *E. coli lptD4213* cells are co-treated with trimethoprim and nisin, the two antibiotics antagonize one another’s activity, as measured by viable cell counts after 2 hours of co-treatment (Figure 6B). Examining the ability to kill MRSA USA300 persister cells yielded an even more striking result in that SCH-79797 could robustly kill these persister cells while the combination of trimethoprim and nisin could not (Figure 6C). These results suggest that the combination of two different antibacterial activities on the same molecular scaffold can, at least in the case of SCH-79797, produce a more potent antibacterial effect than co-treating with two antibiotics with the two separate targeting activities.

### The chemical basis of the two MoAs of SCH-79797

SCH-79797 consists of a pyrroloquinazolinediamine core that is substituted with an isopropylphenyl group on one side and a cyclopropyl moiety on the other. In order to test the function of the pyrroloquinazolinediamine core on the antibiotic activity of SCH-79797, we synthesized a derivative of SCH-79797 (Two-Headed-Monster-10, or THM-10) that lacks both side groups (Figure 7A). When compared to the parent molecule SCH-79797, removing the isopropylphenyl and cyclopropyl groups increased the efficacy against *E. coli lptD4213* but decreased the potency against *B. subtilis* W168, MRSA USA300, and *A. baumannii* AB17978 (Figure 7B). This suggests that while the decorations around the pyrroloquinazolinediamine core are important, they are not strictly necessary for SCH-79797’s antibiotic activity.

**Figure 7.**
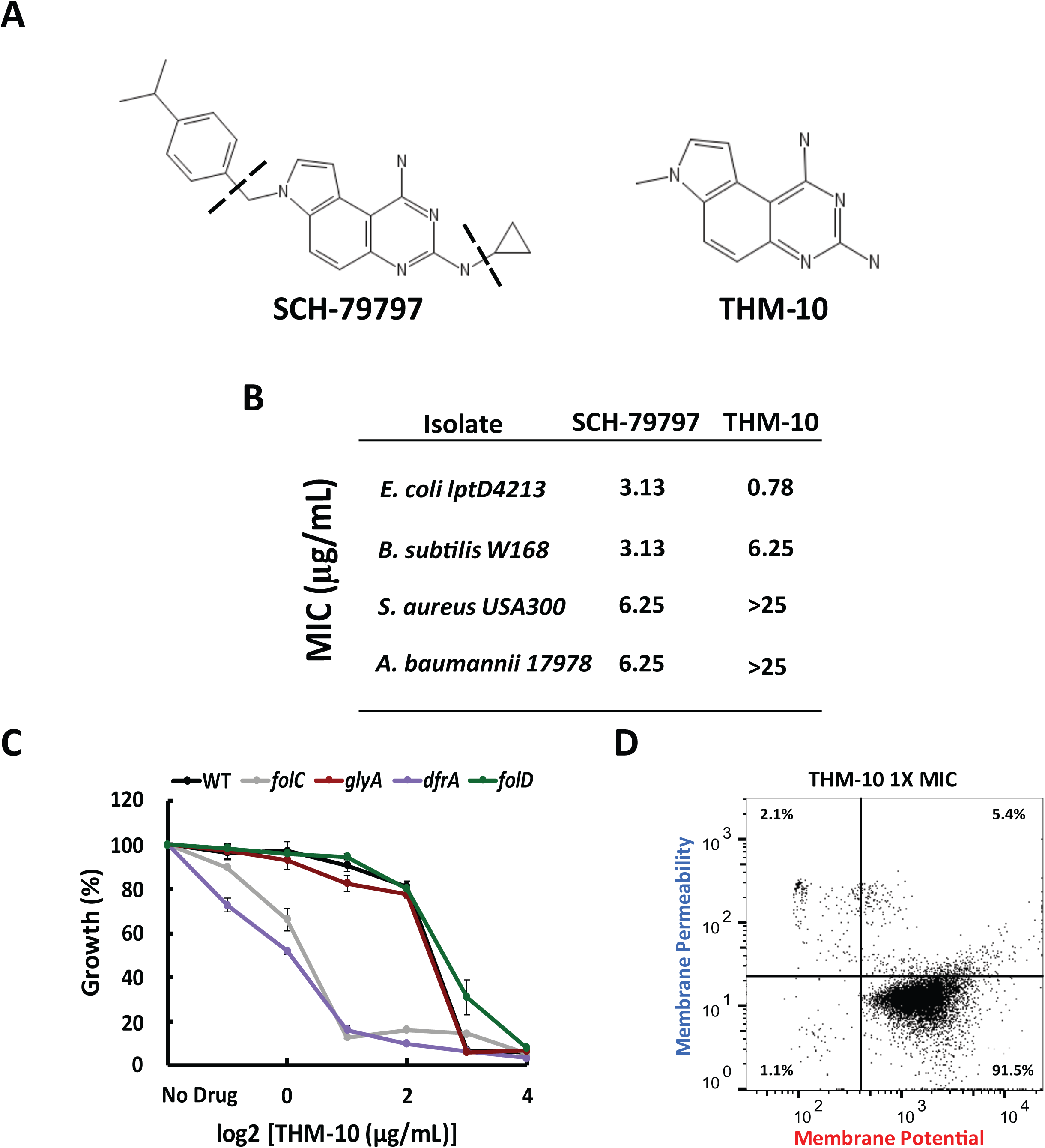
A Derivative of SCH-79797 helps elucidate its two MoAs. A. The structures of SCH-79797 and the pyrroloquinazolinediamine core lacking the side chains, THM-10. B. The MICs of SCH-79797 and THM-10 against *E. coli lptD4213, B. subtilis* W168, *S. aureus* MRSA USA300, and *A. baumannii* AB17978. C. The growth of CRISPRi *B. subtilis* knockdown mutants involved in folate synthesis relative to a DMSO-treated control after THM-10 treatment. Bacterial growth was measured for 14h and the final optical density (OD600) of each condition was plotted against drug concentration. Each data point represents 2 independent replicates. Mean ± s.d. are shown. D. Flow cytometry analysis of the membrane potential and permeability of *E. coli lptD4213* cells after 15 min. incubation with 1X MIC THM-10.

To determine whether the pyrroloquinazolinediamine core of SCH-79797 is specifically involved in targeting folate metabolism or membrane integrity, we assessed the activity of THM-10 using the *dfrA* and *folC* CRISPRi hypersensitivity assay and the quantitative flow cytometry membrane integrity assay. The CRISPRi hypersensitivity assay indicated that THM-10 maintains the ability to inhibit folate metabolism, suggesting that the pyrroloquinazolinediamine core is sufficient to target DHFR (Figure 7C). However, unlike SCH-79797, DIOC_2_(3) and TO-PRO-3 staining showed that THM-10 does not disrupt membrane polarity or permeability (Figure 7D). These findings suggest that the pyrroloquinazolinediamine core of SCH-79797 targets DHFR while the isopropylbenzene and/or cyclopropyl side groups help SCH-79797 disrupt membrane polarization and permeability.

## DISCUSSION

Due to the rise in resistance to known antibiotics, there is an acute need for new antibiotics with the key features of having unique MoAs, potency towards Gram-negatives, and reduced susceptibility to resistance. Here we describe a promising compound, SCH-79797, that is effective in an animal host and addresses these key criteria: it has a unique dual-targeting MoA, kills both Gram-negative and Gram-positive pathogens, and exhibits an undetectably low frequency of resistance. We also describe a systems-level pipeline that combines independent orthogonal approaches to characterize the MoA of SCH-79797 in the absence of resistant mutants. Specifically, we used Bacterial Cytological Profiling (BCP) classification to categorize the MoA of SCH-79797 as distinct from those of 37 known antibiotics (Figure 2B). We then used thermal proteome profiling to identify DHFR as a candidate binding partner of SCH-79797 and confirmed that SCH-79797 inhibits folate metabolism through metabolomic analysis and CRISPRi genetic hypersensitivity (Figure 3B, 4B and C). Finally, we confirmed that SCH-79797 directly inhibits DHFR activity *in vitro* by acting competitively towards its DHF substrate (Figure 4D and E). The BCP images also alerted us to a second potential target for SCH-79797, the bacterial membrane. Quantitative flow cytometry with dyes that report on membrane permeability and polarity confirmed that SCH-79797 has a folate-independent effect on bacterial membrane integrity (Figure 5C). Together, these assays constitute a pipeline that can be used in the future to rapidly characterize antibiotic MoAs *de novo*. Such a pipeline is especially important for compounds such as SCH-79797 that are not prone to resistance and do not mimic known MoAs. BCP, thermal proteome profiling, metabolomics, CRISPRi sensitivity, and flow cytometry are all assays that can be performed in small volumes, such that they can be readily scaled without the need for synthesizing large amounts of the compound in question. We also note that the orthogonal nature of the assays enables the independent identification of multiple MoAs, which may help in the discovery of unique antibiotic classes.

Both of the targets of SCH-79797 are relevant for its function as an antibiotic. The CRISPRi and metabolomic studies demonstrate that SCH-79797 actively disrupts folate metabolism in multiple bacterial species (Figure 4B and C). Meanwhile, the flow cytometry assay demonstrates that SCH-79797 simultaneously disrupts membrane integrity even though folate inhibition itself has no effect on the membrane (Figure 5C). The ability of SCH-79797 to disrupt membrane integrity is particularly interesting given that similar membrane-disruptors like nisin are typically selective for Gram-positive bacteria (Zhou et al., 2016), while SCH-79797 also proved potent against Gram-negative pathogens like *A. baumannii, N. gonorrhoeae*, and pathogenic *E. coli* (Figure 1A). Host toxicity is often a concern for membrane-targeting antibiotics, but SCH-79797 was well tolerated by *G. mellonella* wax worms at 4 times its MIC towards *A. baumannii* (Figure S2A-B) and a recent study of retinoid derivatives provided proof-of-principle that small molecules can preferentially target bacterial membranes (Kim et al., 2018). While SCH-79797 already has relatively low toxicity in a *G. mellonella* model, future biophysical characterization and medicinal chemistry will help to further reduce its toxicity. Similarly, the increased potency of the THM-10 derivative towards *E. coli lptD4213* suggests that medicinal chemistry holds promise for improving the potency of SCH-79797.

The undetectably low frequency of resistance to SCH-79797 could result from its two distinct targets. Specifically, we were successful in isolating resistance mutants for mimics of each of its two individual targets, trimethoprim and nisin, but not for SCH-79797 (Figure 1E). The average mutation rate in *E. coli* is 2.1 × 10^−7^ per gene per generation (Chen and Zhang, 2013). If *E. coli* required 2 mutations to acquire resistance to SCH-79797, the number of bacteria that would be necessary to find a resistant mutant would be in the range of 10^−14^. Humans are estimated to carry roughly 4×10^13^ bacteria in total, so such low frequencies of resistance would be unlikely to result in resistant mutants in a clinical context.

Our studies suggest that SCH-79797 is more potent than combination treatment with antibiotics that mimic its two activities, the DHFR-inhibitor trimethoprim and the membrane-disruptor nisin. Co-treatment with trimethoprim and nisin showed antagonistic interactions (Figure 6B), while MRSA persister cells were killed by SCH-79797 but not by combined treatment with trimethoprim and nisin (Figure 6C). A potential explanation for the potency of SCH-79797 is that recruiting a DHFR inhibitor to the membrane could increase its effective concentration or potentiate its inactivation of DHFR by sequestering it. Permeabilizing the membrane could also enhance the access of SCH-79797 to its cytoplasmic DHFR target. The difference between SCH-79797 and the combination of trimethoprim and nisin could also be based on non-primary target effects such as differences in drug uptake or efflux. Membrane-targeting molecules can act either synergistically or antagonistically with antibiotics with different MoA’s (Brochado et al., 2018). Since trimethoprim and nisin antagonize one another separately but DHFR inhibition and membrane disruption synergize in the context of SCH-79797, combining antibiotic activities onto the same molecule could present a solution for bypassing this antagonistic effect. In any event, our results suggest that despite the promise of combination antibiotic therapies (Brochado et al., 2018; Tyers and Wright, 2019), an even more powerful approach could be to combine different targeting moieties onto the same chemical scaffold.

## MATERIALS AND METHODS

### Bacterial strains and growth conditions

A complete list of strains and growth medias used to grow each bacterium are listed in Supplementary Table 1. The *E. coli* strain, *lptD4213*, was derived from *E. coli* MC4100 and obtained from the lab of Tom Silhavy (Princeton University). Unless otherwise stated, cells were grown from single colonies grown overnight at 37°C in Luria Broth (LB).

### Antibiotics

SCH-79797 dihydrochloride was purchased from Tocris Bioscience. All other antibiotics were purchased from MP Biomedicals at the highest possible purity. All antibiotics except for nisin and gentamicin were dissolved in sterile 100% DMSO. For enzymatic studies, SCH-79797 was dissolved in 100% EtOH since DMSO is toxic to DHFR protein. Nisin was dissolved in sterile 0.02N HCl and gentamicin was dissolved in sterile deionized water. The minimum inhibitory concentration of each antibiotic was defined as the lowest concentration of antibiotic that resulted in no visible growth. MICs were measured using 2-fold dilutions of each antibiotic in 96-well plates and cell growth was monitored by measuring the OD600.

### Compound library

Compounds were sourced from commercial vendors: MicrosourceDiversity, Aldrich, Sellekchem, Chiromics, and Chembridge. Each compound was screened for antibiotic activity against *E. coli lptD4213* at 50μM in DMSO. *E. coli lptD4213* were grown in Terrific Broth. After normalizing for plate-to-plate variation, we used an OD600 of half the median plate OD600 as our cutoff, below which any compound was assumed to have inhibited the growth of *E. coli lptD4213* and above which compounds were assumed to be ineffective. Compounds that either had not been previously identified as antibiotics or had unknown or ambiguous MoAs were chosen for further investigation and their MIC’s were measured using the microdilution method described above.

### *Galleria mellonella* killing assay

All *Galleria mellonella* larvae were obtained from Vita-Bugs©, distributed through PetCo© (San Diego, CA), and kept in a 20°C chamber. All injections were administered using a sterile 1 ml syringe attached to a KD Scientific pump delivered at a rate of 250 µl/min to the fourth leg of the worm, which was sterilized with EtOH. *A. baumannii* AB17978 (10^5^ CFU/larva) and drug dissolved in DMSO (SCH-79797 at 66.6 μg/larva, gentamicin at 6 μg/larva, rifampicin at 66.6 μg/larva) were pre-mixed prior to injection. The viability of each injected larva was determined by prodding each larva with a dowel and observing whether there was subsequent movement.

### Colony Forming Units Assay

Overnight *E. coli lptD4213* or *S. aureus* MRSA USA300 cultures were diluted 1:100 in fresh media and grown to early-mid exponential phase (OD600 = 0.4-0.6). Each culture was then diluted 1:10 into fresh media and then treated with the desired concentration of each antibiotic. Each time point was taken by removing 150μL from each treatment condition and diluting 1:10. 6 dilutions of each condition were then plated in the absence of antibiotic and grown at 37°C overnight. CFU’s were measured by counting the resulting number of colonies the next day.

### Serial passaging Assay to determine the dynamics of resistance emergence

2 independent overnight cultures for each of S. aureus MRSA USA300 and *A. baumannii* AB17978 were diluted 1:150 in LB and treated with 2-fold dilutions of each antibiotic. Bacteria were grown for 14h in a 96-well plate in LB at 37°C and bacterial growth was measured by monitoring changed in OD600 in a Tecan microplate reader. Cultures that grew in 0.5X MIC of each antibiotic were streaked out onto plain LB agar plates and single colonies were used to create the inoculum of the next passage and 750μL of each inoculum was stored in 25% glycerol at −80°C. Confirmation of resistance to each antibiotic was determined by re-measuring the MICs of each glycerol freezer stock.

### Bacterial Cytological Profiling

Experiments described in Figure 2 were performed as described in (Nonejuie et al., 2013). In later experiments (Figure 6A), overnight *E. coli lptD4213* cultures were diluted 1:100 and grown to early-mid exponential phase (OD600 = 0.4-0.6). Each culture was then diluted 1:10 into fresh LB and treated with the desired concentration of antibiotic for 10 minutes. Following antibiotic treatment, cells were stained with 0.5 μM SYTOX Green, 1μg/mL FM4-64, and 2 μg/mL DAPI. Each stained culture was then spotted onto a 1.5% agar pad supplemented with casamino acids and 20% glucose in M63. The *E. coli* cells were segmented, and single-cell features were extracted using a custom Matlab code. Principal component analysis was performed using the prcomp function in R and clustering was performed using the single-linkage method.

### Thermal Proteome Profiling assay

Thermal proteome profiling experiments were performed as described by (Mateus et al., 2018) Briefly, whole cell samples were treated with 0.6, 1.1, 2.2, 4.4μg/mL SCH-79797 and 0.1, 0.5, 2.3, 11.6μg/mL trimethoprim. Cell lysate samples were treated with 0.4, 1.8, 8.9, 44.4μg/mL SCH-79797 and 0.1, 0.5, 2.3, 11.6μg/mL trimethoprim. The mass spectrometry proteomics data have been deposited to the ProteomeXchange Consortium via the PRIDE partner repository with the dataset identifier PXD013673. After heat treatment, the soluble fraction was collected, digested with trypsin and peptides were labeled with tandem mass tags (Werner et al., 2014). Samples were subjected to two-dimensional liquid chromatography and analyzed on a Q Exactive Plus mass spectrometer (Thermo Fisher Scientific). While the data collected is proteome wide, three different cutoffs were used to define three classes of potential targets. The color of the point indicates the signal maximal effect size across all temperatures and the largest change in abundance across all concentrations was selected for continued analysis (Dots, Figure 3.) A mild effect indicates at least one temperature had a change in abundance of at least 25% in both whole cell and cell lysate treatments (Triangles, Figure 3) Proteins that had a change in abundance of at least 25% at three or more temperatures (Squares, Figure 3). To be considered a consistent effect, the change in abundance of the protein had to show the same sign at least 90% of the time and have an effect size of at least 1 (2-fold) in either whole cells or cell lysates.

### *B. subtilis* CRISPRi hypersensitivity assay

We used the indicated mutants from the CRISPR interference (CRISPRi) library as curated by (Peters et al., 2016) to measure the sensitivity of mutants involved in folate synthesis to SCH-79797. MICs were measured using 2-fold dilutions of each antibiotic in 96-well plates and cell growth was monitored by measuring the OD600 in a Tecan microplate reader.

### Metabolomics

Overnight *E. coli* NCM3722 cultures were grown and diluted 1:100 in Gutnick Minimal Media and grown to early-mid exponential phase (OD600 = 0.4-0.6). Cultures were treated with either 1X MIC SCH-79797 (13.89μg/mL) or 1X MIC trimethoprim (0.15μg/mL) for 15 minutes. Folates were extracted by vacuum filtering 15mL of treated cells using 0.45μm HNWP Millipore nylon membranes and immediately placing filters into an ice-cold quenching solution containing 40:40:20 MetOH:acetonitrile:25Mm NH_4_OAc + 0.1% sodium Ascorbic in HPLC H_2_O. The resulting solution was then centrifuged at 16,000×g for 1.5 min at 4°C and the supernatant was saved for mass spectrometry analysis which was performed as described in (Chen et al., 2017)

### DHFR Enzymatic Activity assay

*E. coli* DHFR enzyme (FolA) was custom purified by Genscript (Piscataway, NJ). The enzymatic activity of DHFR with and without SCH-79797 treatment was measured using the Dihydrofolate Reductase Assay kit from Sigma. Briefly, DHFR activity was measured by monitoring the change in sample absorbance at 340nm due to DHFR-dependent NADPH consumption.

### Persister cell assay

Stationary phase *S. aureus* cells have the same antibiotic tolerant properties of persister cells (Kim et al., 2018). Thus, overnight MRSA USA300 cultures were used to measure the effectiveness of SCH-79797 against persister cells. An overnight *S. aureus* MRSA USA300 culture was diluted 1:100 in PBS prior and then cell viability was determined using the same CFU analysis described above.

### Membrane potential and permeability assay

The BacLight Bacterial Membrane Potential kit from Sigma was used to measure the effect of SCH-79797 treatment on bacterial membrane potential. This kit uses DiOC2(3) to measure a cell’s membrane potential. This dye concentrates in cells with an active membrane potential causing the emission of DiOC2(3) to shift from green (488nm excitation, 525/50nm bandpass filter for emission) to red (488nm ex, 610/20 bp filter). As a result, by measuring the fluorescence shift from red to green, we can detect changes in a cell’s membrane potential. TO-PRO-3 is a dye that is excluded from cells with a healthy membrane. Thus, we can detect membrane damage by measuring the far-red fluorescence intensity (640nm ex, 670/30nm bp filter). The LSRII flow cytometer (BD Biosciences) at the Flow Cytometry Resource Facility, Princeton University, was used to measure the fluorescent intensities of both dyes in response to antibiotic treatment. Data was analyzed using FlowJo v10 software as described in (Novo et al., 1999).

