## Supplementary Figures and Legends for "A dual-mechanism antibiotic targets Gram-negative bacteria and avoids drug resistance"

### Supplementary Materials

#### Supplementary Table 1: Bacterial Strains and Growth Media

| Strain | Growth Media |
|---|---|
| *B. cepecia* | Cation-adjusted Mueller-Hinton Broth II |
| *E. coli WT BW25113* | Luria Broth |
| *E. coli CFT073* | Luria Broth |
| *C. difficile* | Reinforced Clostridial Medium |
| *P. acnes* | Reinforced Clostridial Medium |
| *H. influenza* | Cation-adjusted Mueller-Hinton Broth II |
| *V. cholerae* | Luria Broth |
| *A. baumannii AB17978* | Luria Broth |
| *A. baumannii AB5075* | Luria Broth |
| *E. coli UPEC CFT073* | Gutnick Minimal Media |
| *E. coli UPEC J96* | Gutnick Minimal Media |
| *E. coli BW25113* | Gutnick Minimal Media |
| *E. coli NCM3722* | Gutnick Minimal Media |
| *S. aureus MRSA COL* | Luria Broth |
| *S. aureus MRSA USA300* | Luria Broth |
| *M. fortuitum* | Cation-adjusted Mueller-Hinton Broth II |
| *N. gonorrhoeae* | Cation-adjusted Mueller-Hinton Broth II |
| *S. aureus VanA* | Cation-adjusted Mueller-Hinton Broth II |
| *S. aureus VISA* | Cation-adjusted Mueller-Hinton Broth II |
| *S. aureus MRSE* | Cation-adjusted Mueller-Hinton Broth II |
| *S. epidermidis* | Cation-adjusted Mueller-Hinton Broth II |
| *B. subtilis W168* | Luria Broth |
| *E. coli lptD4213* | Luria Broth |
| *E. faecium* | Cation-adjusted Mueller-Hinton Broth II |

| | |
|---|---|
| *S. pneumoniae* | Cation-adjusted Muller-Hinton Broth II with 5% lysed horse blood |

**Supplementary Table 2: The 14 features evaluated in BCP analysis**

| **Features of BCP Analysis** |
|---|
| Cell Area |
| Cell Length |
| Cell Width |
| Cell Eccentricity |
| Cell Perimeter |
| Nucleoid Area |
| Ratio of Nucleoid to Cell |
| Nucleoid Eccentricity |
| DNA Length |
| DNA Width |
| DNA Perimeter |
| Mean Sytox intensity |
| Mean FM4-64 intensity |
| Mean Dapi intensity |

#### Supplementary Figure Legends

**Figure S1. SCH-79797 is bactericidal against *S. aureus*.** Colony forming units (CFU $ml^{-1}$) after 3-hour treatment of *S. aureus* MRSA USA300 with 1% DMSO, 1X MIC SCH-79797 and 5X MIC novobiocin. Each data point represents 3 independent samples and 3 technical replicates. Mean ± s.d. are shown.

**Figure S2. SCH-79797 is an effective antibiotic in an infection model of *G. mellonella* by *A. baumannii.*** A-B. The percent survival of non-infected *G. mellonella* wax worms after treatment with 2µl/larva of 100% DMSO, 67µg/larva SCH-79797, 6µg/larva gentamicin, and

67μg/larva rifampicin. Data in (A) represents a typical cohort (n = 12) from a biological triplicate and the pooled results are presented in (B). Mantel-cox statistics for the cohort were calculated with PRISM. C. The percent survival of *A. baumannii* infected *G. mellonella* wax worms after treatment with 67μg/larva SCH-79797, 6μg/larva gentamicin, and 67μg/larva rifampicin. Data represents the pooled results from a biological triplicate. D. The average survival of drug-treated *A. baumannii* infected *G. mellonella* wax worms relative to larvae treated with DMSO. Data represents the pooled results from a biological triplicate.

**Figure S3. SCH-79797 is not prone to resistance in both *S. aureus* and *A. baumannii*.** A. Fold increase in resistance of *S. aureus* MRSA USA300 to SCH-79797, novobiocin, trimethoprim, and nisin after 25 days of serial passaging in 0.5X MIC of each drug and plotted on a log2 scale. Resistance was confirmed by remeasuring MIC's from aliquots of each passage that were collected and stored at -80$^{o}$C. B. Fold increase in resistance of *A. baumannii* AB17978 to SCH-79797 and gentamicin after 5 days of serial passaging in 0.5X MIC of each drug and plotted on a log2 scale. Resistance was confirmed as described above. The experiment was performed in duplicate and identical results were found in both cases.

**Figure S4. Thermal stability of DHFR increases after SCH-79797 and Trimethoprim treatment.** A-B. The relative thermal stability of DHFR after treatment of whole cell and cell lysate samples with (A) SCH-79797 and (B) trimethoprim. Changes in thermal stability were determined by measuring changes in the abundance of DHFR across 10 different temperatures ranging from 42-72$^{o}$C and 4 drug concentrations and a vehicle control.

**Figure S5. CRISPRi mutants not involved in folate metabolism are not sensitized to SCH-79797.** A. The growth of CRISPRi *B. subtilis* knockdown mutants relative to a DMSO-treated control after SCH-79797 treatment. Bacterial growth was measured for 14h and the final optical density (OD600) of each condition was plotted against drug concentration. Each data point represents 2 independent replicates. Mean ± s.d. are shown.

**Figure S6. Treatment with ampicillin, rifampicin, and novobiocin does not disrupt membrane integrity.** A. Flow cytometry analysis of the membrane potential and permeability of *E. coli* lptD4213 cells after 15 min. incubation with 2X MIC ampicillin, rifampicin, novobiocin.

**Figure S7. SCH-79797 disrupts *B. subtilis* W168 membrane integrity.** A. Flow cytometry analysis of the membrane potential and permeability of *B. subtilis* W168 cells after 15 min. incubation with 1% DMSO, 1X MIC SCH-79797, 2X MIC SCH-79797.

#### Supplementary Figure 1

**A**

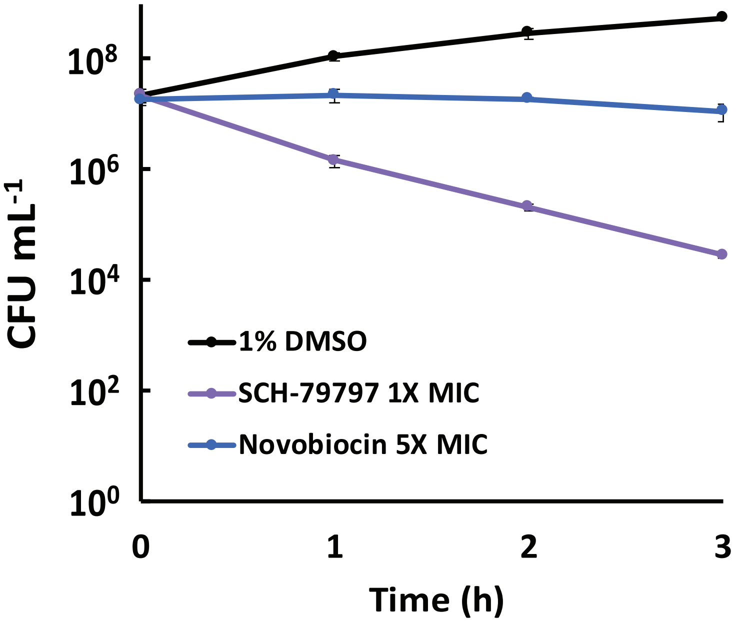

### Supplementary Figure 2

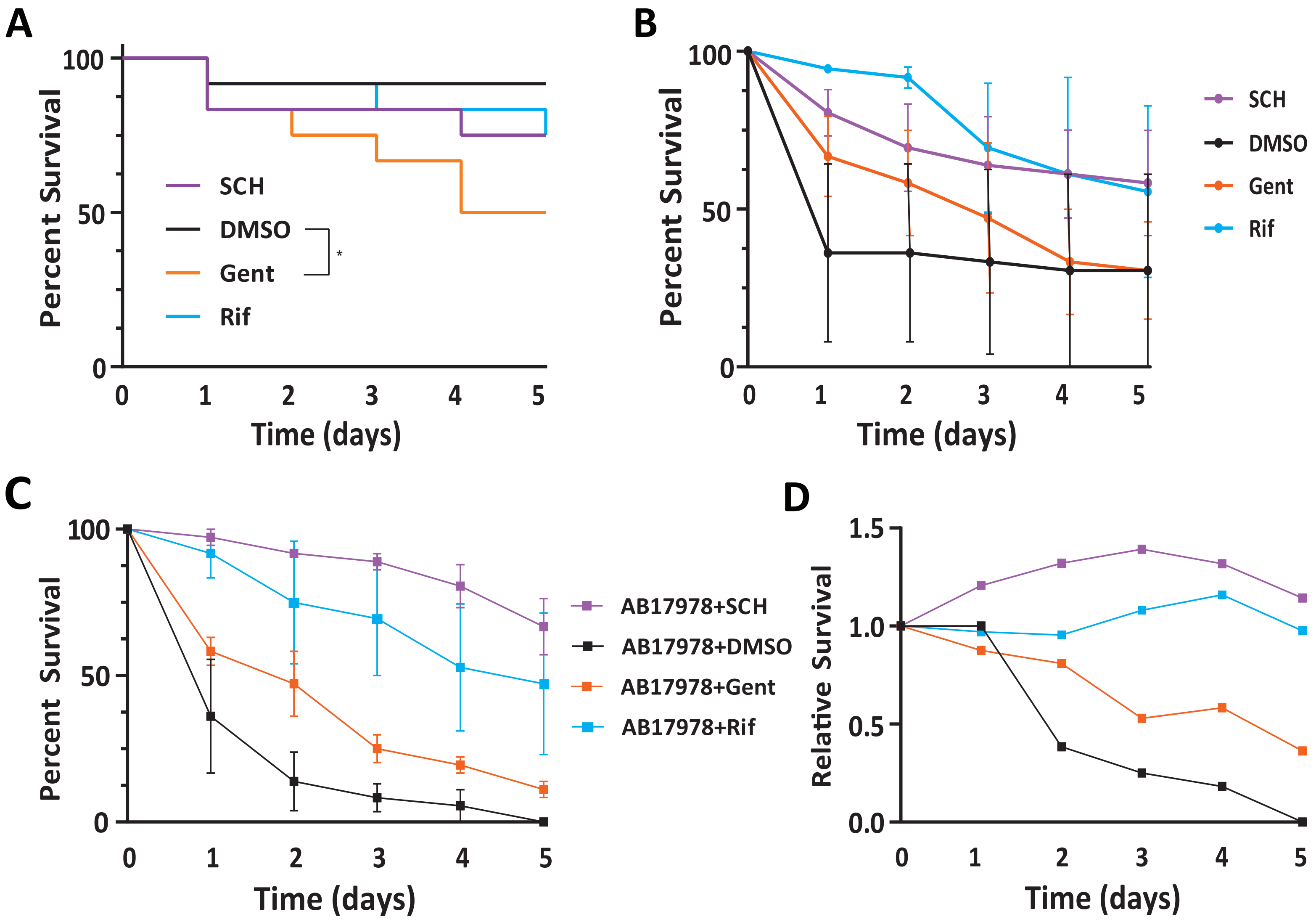

#### Supplementary Figure 3

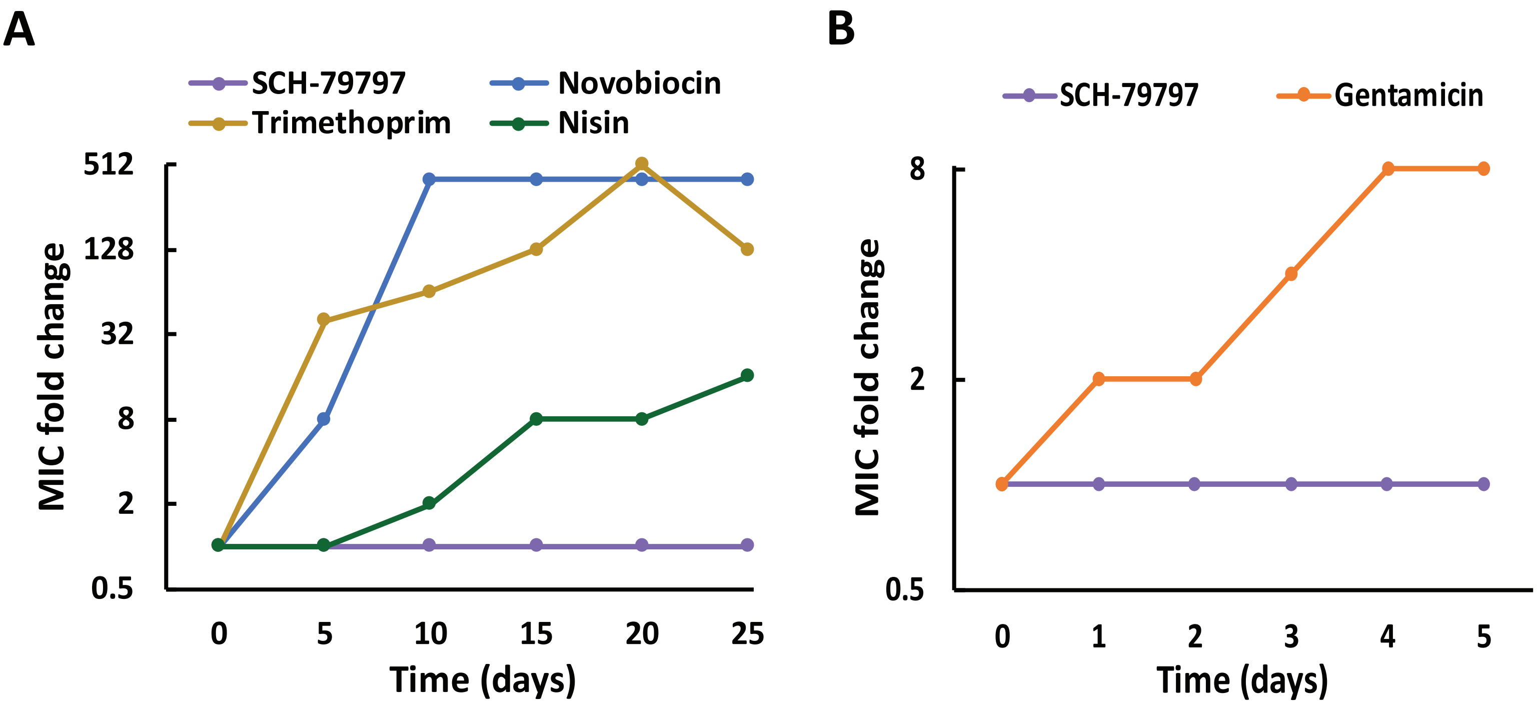

### Supplementary Figure 4

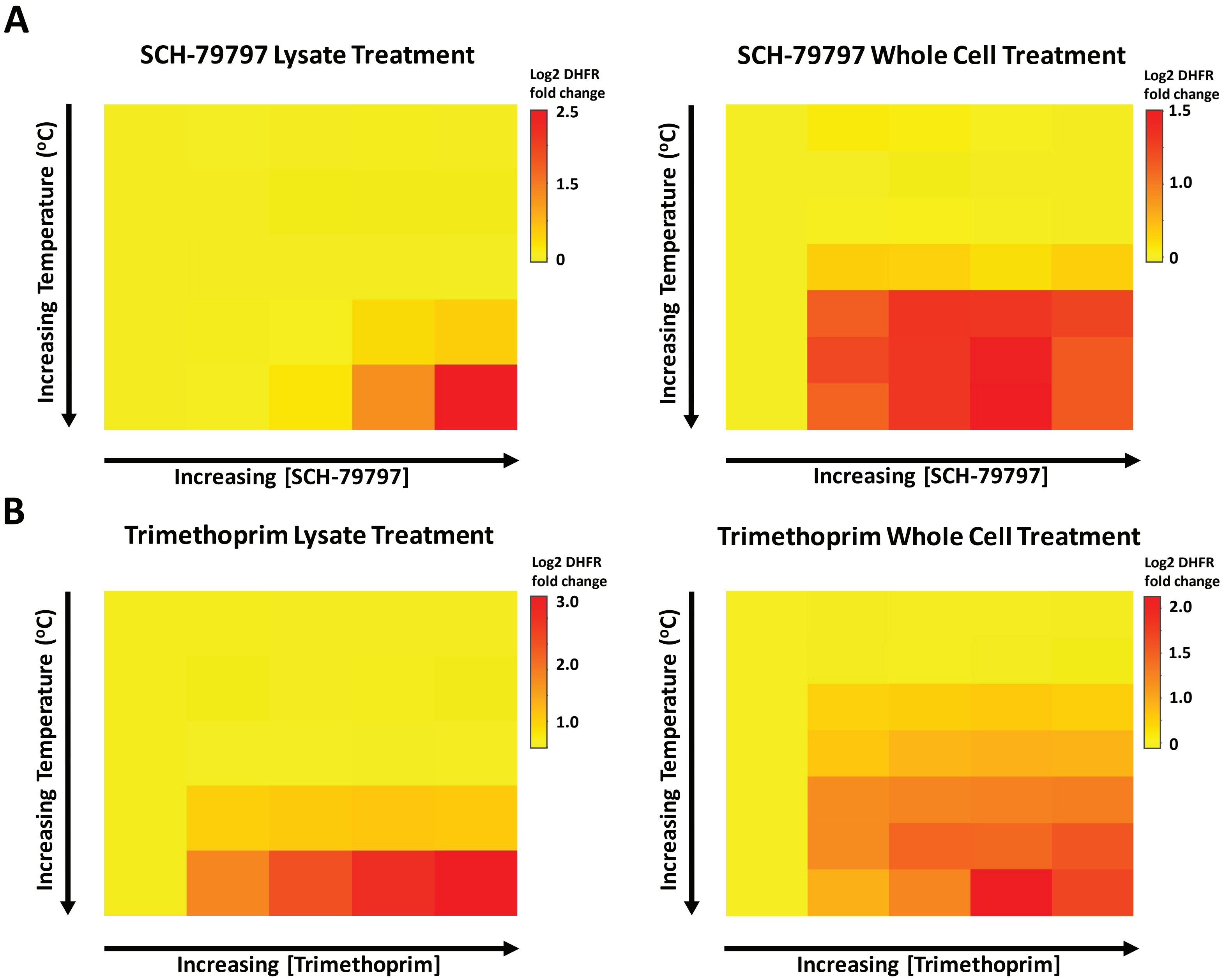

### Supplementary Figure 5

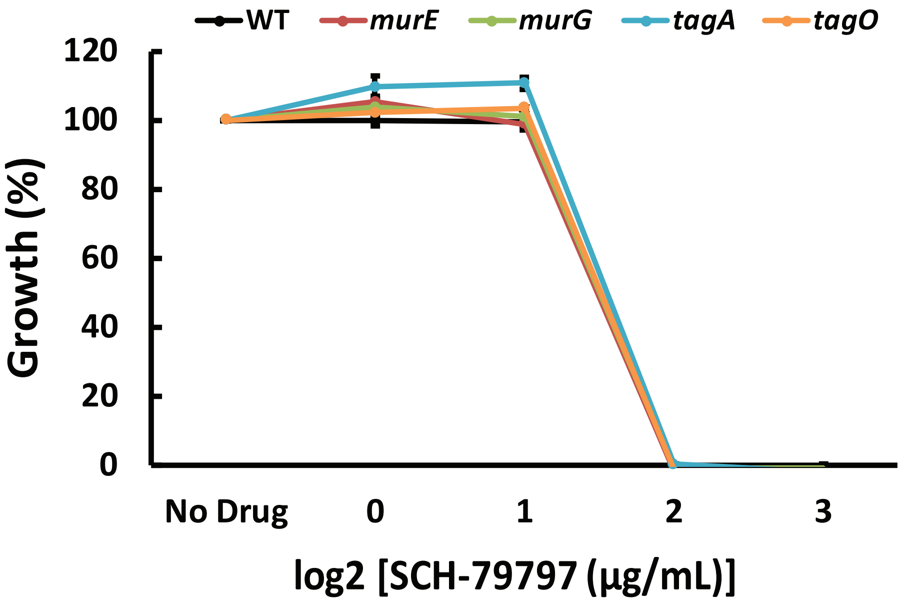

#### Supplementary Figure 6

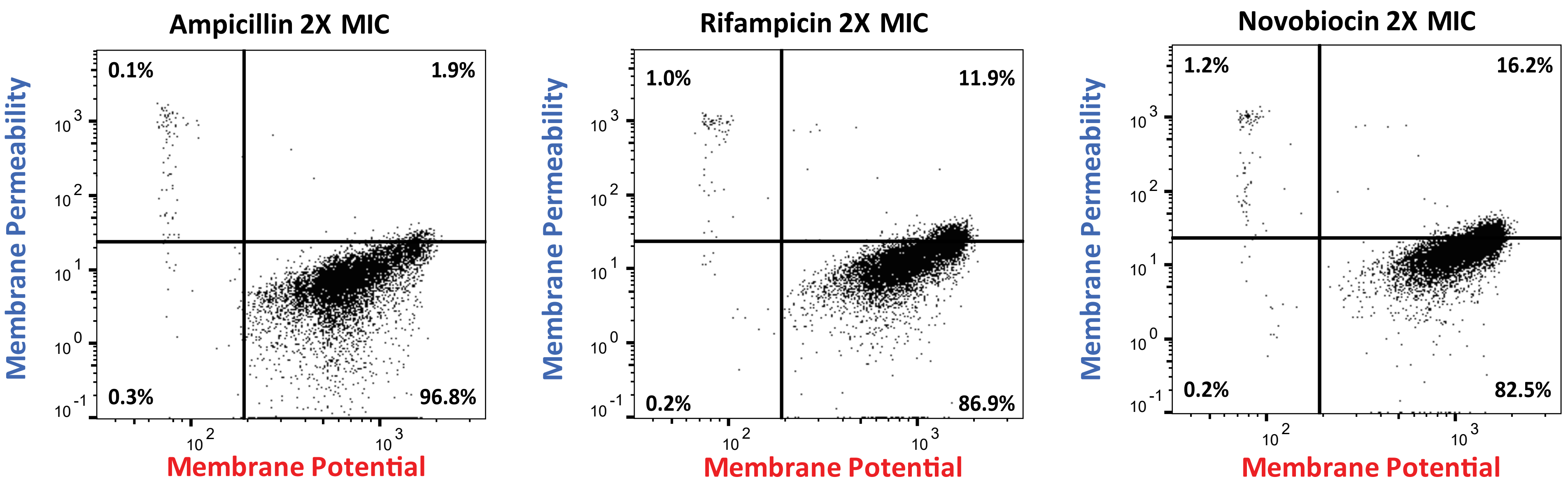

### Supplementary Figure 7

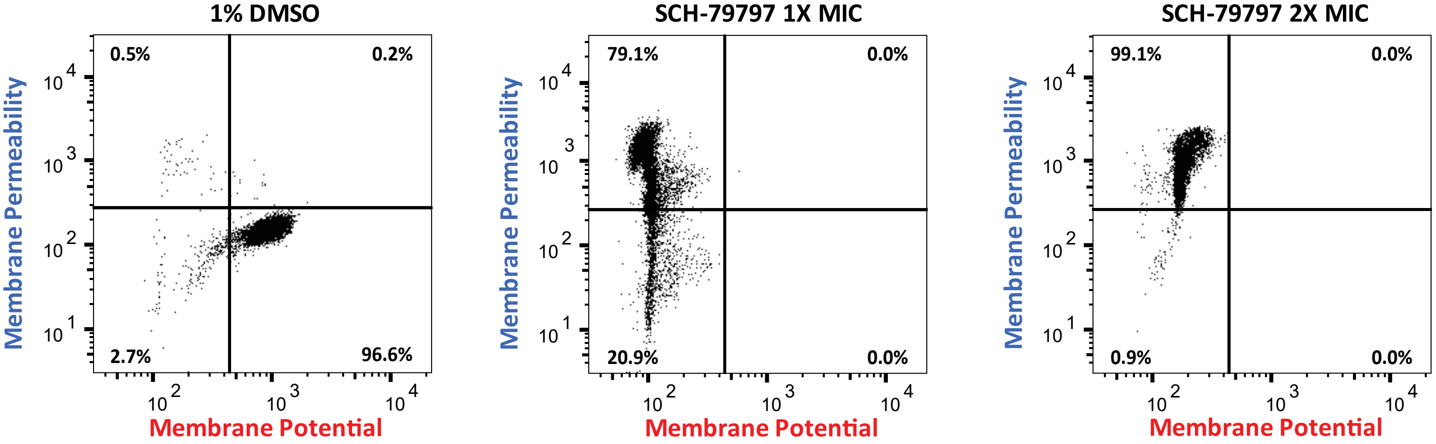
